# Neural processing of rhythmic speech by children with developmental language disorder (DLD): An EEG study

**DOI:** 10.1101/2024.03.29.587020

**Authors:** Mahmoud Keshavarzi, Susan Richards, Georgia Feltham, Lyla Parvez, Usha Goswami

## Abstract

Sensitivity to rhythmic and prosodic cues in speech has been described as a precursor of language acquisition. Consequently, atypical rhythmic processing during infancy and early childhood has been considered a risk factor for developmental language disorders. Despite many behavioural studies, the neural processing of rhythmic speech has not yet been explored in children with developmental language disorder (DLD). Here we utilise EEG to investigate the neural processing of rhythmic speech by 9-year-old children with and without DLD. In the current study, we investigate phase entrainment, angular velocity, power, event related potentials (ERPs), phase-amplitude coupling (PAC) and phase-phase coupling (PPC), at three frequency bands selected on the basis of the prior literature, delta, theta and low gamma. We predicted a different phase of entrainment in the delta band in children with DLD, and also greater theta power, atypical cross-frequency coupling and possibly atypical gamma-band responses. Contrary to prediction, children with DLD demonstrated significant and equivalent phase entrainment in the delta and theta bands to control children. However, only the control children showed significant phase entrainment in the low gamma band. The children with DLD also exhibited significantly more theta and low gamma power compared to the control children, and there was a significant gamma-band difference in angular velocity between the two groups. Finally, group resultant phase analyses showed that low-frequency phase (delta and theta) affected gamma oscillations differently by group. These EEG data show important differences between children with and without DLD in the neural mechanisms underpinning the processing of rhythmic speech. The findings are discussed in terms of auditory theories of DLD, particularly Temporal Sampling theory.

## 1. Introduction

A growing number of cognitive behavioural studies have investigated rhythmic and/or musical processing in children with developmental language disorder (DLD), and have shown that atypical rhythm processing is associated with speech and language processing problems (Bedoin et al., 2016; Corriveau et al., 2007; Corriveau & Goswami, 2009; Cumming et al., 2015a,b; Przybylski et al., 2013; Richards & Goswami, 2015, 2019; Sallat & Jentschke, 2015). This apparent link between rhythm processing and language processing has aroused considerable interest given its potential implications for remediation. For example, following a comprehensive review of the literature, Ladányi et al. (2020) proposed the Atypical Rhythm Risk (ARR) hypothesis, suggesting that individuals with atypical rhythm processing are at higher risk for developmental speech/language disorders including developmental dyslexia, DLD and stuttering. DLD is the focus of the current study, and affects between 3 – 7% of children in all cultures (Bishop et al., 2017). The clinical diagnosis for such children was previously Specific Language Impairment, however the multinational and multidisciplinary CATALISE study recommended a broadening of diagnostic features and a change in terminology to DLD (Bishop et al., 2017). Children given a diagnosis of DLD have a range of language difficulties that cause functional impairments in their daily lives, yet have no known biomedical aetiology. The core dimensions of difficulty are problems with syntax and grammar, problems with word semantics, impairments in verbal memory and problems with phonology (see Bishop, 2013; Leonard, 2014). The possibility that atypical *neural* processing of rhythm is associated with DLD has not yet been investigated, and is the focus of the current study.

At the sensory/neural level, the ARR hypothesis is captured by Temporal Sampling (TS) theory (Goswami, 2011; 2015, 2022). TS theory was originally proposed to explain the rhythmic impairments found in children with developmental dyslexia, a disorder of written language processing that presents with core phonological impairments (Snowling, 2000; Stanovich, 1988; Swan & Goswami, 1997). However, prior TS-driven sensory and behavioural studies of rhythmic processing by children with dyslexia and children with DLD have shown many shared features between children diagnosed with these distinct disorders (Goswami, 2022, for review). For example, both developmental disorders exhibit impaired sensory discrimination of amplitude rise times (important acoustic cues to rhythm, Beattie & Manis, 2012; Corriveau et al., 2007; Corriveau & Goswami, 2009; Cumming et al., 2015a; Fraser et al., 2010; Goswami et al., 2002, 2020; Richards & Goswami, 2015, 2019) and relative insensitivity to syllable stress patterns in words and phrases (Caccia & LoRusso, 2019a,b; Cumming et al., 2015a,b; Goswami et al., 2010, 2013; Richards & Goswami, 2015, 2019). To explain these sensory and cognitive difficulties with rhythm, TS theory suggests that the neural processing of rhythm is impaired in affected children. The TS framework (TSF) proposes that the oscillatory encoding of slower amplitude modulation (AM) information in the speech amplitude envelope (AE) is impaired in children with DLD and dyslexia, particularly at modulation rates <10 Hz that are known to carry information about speech rhythm and syllable stress patterns (Greenberg, 2006). The AE is the slow-varying energy contour of speech, and contains a range of AM patterns, created by hierarchical nesting of AMs at different temporal rates (Ghitza & Greenberg, 2009; Goswami & Leong, 2013; Leong & Goswami, 2015). Computational speech modelling studies show that the phase relations between these different AM rates (delta-theta, theta-beta/low gamma) provide systematic statistical cues to both phonological units such as stressed vs unstressed syllables, syllables, and onset-rimes (acoustic-emergent phonology, Leong & Goswami, 2015) and also to aspects of grammar such as plural marking (Flanagan & Goswami, 2018). By TS theory, therefore, impaired oscillatory encoding (cortical tracking) of one or more of the AM rates nested in the AE, or of the phase-phase relations between different AM bands, could cause atypical development of one or more aspects of language.

The adult speech processing literature has shown that the speech AE is tracked by endogenous oscillations in a number of electrophysiological bands, the most important being delta (0.5-4 Hz), theta (4-8 Hz) and low gamma (∼30Hz, Giraud & Poeppel, 2012; Gross et al., 2013). The sensory trigger for cortical tracking at these different temporal rates is amplitude rise time (Doelling et al., 2014; Gross et al., 2013). These different temporal processing rates are proposed to support the parsing of different levels of phonological information in the speech signal (Giraud & Poeppel, 2012; Poeppel, 2014). Crucial information for successful linguistic decoding is carried at multiple temporal scales, including intonation-level or prosodic information at the scale of 500–2000 ms (argued to be encoded by delta band responses), syllabic information that is correlated to the acoustic envelope of speech at the scale of 150– 300 ms (encoded by theta band responses), and rapidly-changing phonetic featural information at the scale of 20–80 ms (encoded by gamma band responses; Giraud & Poeppel, 2012). Speech encoding is shown to be supported by the sliding and resetting of neural temporal windows that match information in the signal to oscillatory responses, implemented as phase locking to key features of the input via phase-resetting of the intrinsic oscillations by amplitude rise times (Poeppel, 2014). Although research with children and infants has shown that phonetic feature information is also encoded by the slower delta and theta band responses (see Di Liberto et al., 2018, 2023), the idea that gamma band responses encode phoneme-relevant information has been very influential in the neural speech encoding literature.

In the current study, children’s neural processing of rhythm was investigated using the syllable repetition task (Power et al., 2012), a task that has previously been administered to both children with and without dyslexia and to infants (Keshavarzi et al., 2022; Ni Choisdealbha et al., 2022, 2023; Power et al., 2013). In this task, EEG is recorded while participants listen to the speech syllable “ba” or “ta” presented audio-visually at a repetition rate of 2 Hz. Both 9-year-old (Keshavarzi et al., 2022) and 13-year-old (Power et al., 2013) children with dyslexia showed atypical delta-band phase synchronisation in the syllable repetition task, exhibiting a different preferred phase to children without dyslexia for the delta band. The theta band was not affected. In the infant studies, which tracked a cohort of 122 infants longitudinally for over 3 years, delta band phase from 2 months of age in the rhythmic syllable repetition task predicted later language outcomes at 12, 18 and 24 months of age (Ni Choisdealbha et al., 2022, 2023). These converging delta-band data from children and infants suggest that there is an optimal (or preferred) phase of entrainment for the accurate and efficient processing of rhythmic inputs, and that individual differences in preferred delta phase have consequences for language outcomes. To date, the rhythmic syllable repetition task has not been administered to children with DLD. That is our aim here. *A priori*, given the similar sensory and cognitive profiles of children with DLD and children with dyslexia in behavioural speech rhythm tasks, we expected that phase entrainment in the delta and theta bands would be significant for both typically-developing (TD) age-matched control and DLD groups, and children with DLD would show a different preferred phase in the delta band compared to TD children (H1).

Prior neural studies of children with DLD have not to date employed rhythmic processing tasks nor investigated the TS framework. Indeed, prior neural DLD studies have instead primarily utilised fMRI and have sought structural and functional differences that may throw light on the aetiology of this developmental disorder. For example, there has been interest in whether hemispheric asymmetries during linguistic processing are different when children with DLD are compared to TD children (Bishop, 2013). While laterality per se has not proved conclusive (Parker et al., 2022), one of the most consistent findings in the functional fMRI literature with DLD children is that left frontal and temporal cortical areas show reduced activation during language processing (Asaridou & Watkins, 2022, for review). Prior MEG studies of DLD using speech processing tasks are consistent with the functional fMRI literature in showing atypical left hemisphere temporal function (Helenius et al., 2009, 2014). Children with DLD also show microstructural differences in dorsal anatomical pathways that connect frontal and temporal areas in fMRI studies, as well as structurally-atypical basal ganglia (Badcock et al., 2012; Lee et al., 2013; Watkins et al., 2002). Although both the temporal areas and the basal ganglia are important for speech processing, speech motor learning and temporal prediction (see Kotz & Schmidt-Kassow, 2015), fMRI studies to date have not revealed which neural mechanisms associated with these areas could be affected in DLD.

The absence of neural rhythm studies with DLD participants is surprising, as adult neural studies show that temporal prediction of the location of stressed syllables facilitates syntactic processing (Kotz & Schmidt-Kassow, 2015). Linguistic analyses show that adults produce a stressed syllable on average every 500 ms, a “beat rate” of 2Hz (Dauer, 1983), a finding that underpins the choice of the 2 Hz repetition rate in the current task. If neural temporal prediction of stressed syllable placement is important developmentally for language acquisition, then it might also be expected *a priori* that dynamic relations (cross-frequency coupling) between the electrophysiological delta and theta bands may be atypical in children with DLD. In the infant speech rhythm studies (Ni Choisdealbha, 2022, 2023), nursery rhymes were also sung to the participating infants while EEG was recorded (Attaheri et al., 2022; Keshavarzi et al., 2024b). While the accuracy of delta band cortical tracking predicted better language outcomes, greater theta power and a higher excitation/inhibition ratio between theta and delta predicted worse language outcomes (Attaheri et al., preprint). Meanwhile an EEG study using a story listening task found that children with DLD had significantly greater delta-theta phase-amplitude coupling (PAC) compared to TD children (Araújo et al., 2024). Accordingly, two further predictions were derived. H2 was that delta-theta cross-frequency coupling may differ between the DLD and TD children in the current study, and H3 was that differences in theta power may be found, with greater theta power for children with DLD.

Nevertheless, most prior EEG studies with DLD children have not adopted a rhythmic nor a speech tracking focus, hence H2 and H3 should be considered tentative only.

Indeed, many of the prior DLD child EEG studies concern the overlap between DLD diagnoses and epileptic seizures (e.g. Mehta et al., 2015; Shafer et al., 2001; Systad et al., 2019). The exception is a series of EEG studies with German-learning infants, which have shown that those at family risk for DLD show impaired neural ERP (event related potential) markers of language learning, such as a delayed N400 (thought to index semantic integration, Friedrich & Friederici, 2005, 2006). Meanwhile, an alternative auditory theory of DLD to TS theory, the rapid auditory processing (RAP) theory (Tallal, 1980, 2004), has proposed that basic auditory processing of brief, rapidly successive acoustic changes is impaired in children with DLD, expected to affect the phoneme level of speech processing (Tallal & Piercy, 1973). Tallal and Piercy argued that phonetic discrimination required temporal processing at rapid timescales of around 40 ms, which would correspond to oscillatory gamma band information. Consistent with RAP, a nonspeech MEG study with DLD children indicated altered temporal organization of gamma-band oscillatory activity when children with DLD processed rapid tone sequences (nonspeech input, Heim et al., 2011). Children listened to pairs of tones separated by 70 ms, and judged the pitch of the second tone. Heim et al. reported significantly reduced amplitude and phase-locking of early (45–75 ms) oscillations in the gamma-band range (29–52 Hz) for the second stimulus. ERP studies with English-learning infants at family risk for DLD using a non-speech tone pair task have shown that at-risk infants have ERPs with larger amplitudes than TD infants when the tones follow each other rapidly in time (gap of 70 ms, Benasich et al., 2006), and also show a longer latency (in the N250) which is right-lateralized. Some studies of at-risk infants also report differences in resting state gamma power compared to TD infants (Ortiz-Mantilla & Benasich, 2013), and in some studies these ERP differences in the gamma band predicted later language skills (English, language measured at 3, 4 and 5 years, Benasich et al., 2008; Choudhury & Benasich, 2011; Gou et al., 2011; Italian, language measured at 20 months, Cantiani et al., 2016, 2019). Accordingly, a final tentative prediction was that gamma band responding could be atypical in the current task for children with DLD (H4).

A gamma band difference between groups would also be consistent with a prior study of rhythmic speech processing that used a modulation band manipulation with children with DLD (Goswami et al., 2016). Goswami et al. (2016) gave children a nursery rhyme recognition task based on speech that was either low-pass filtered (< 4 Hz, providing delta band information only) or band-pass filtered (∼33Hz, providing faster low gamma band modulations only). In both conditions, the speech input was degraded and difficult to recognise. The children’s task was to recognise the matching nursery rhyme by making a selection in a picture-based multiple choice format. Goswami et al. reported that 9-year-old children with DLD were worse than TD age-matched children when selecting bandpass-filtered targets, but not when selecting low-pass filtered targets, possibly suggestive of preserved delta band processing. Children with both DLD and dyslexia (DLD children with phonological impairments) were worse than TD children at recognising the nursery rhymes in both filtered speech conditions. All groups of children found it easier to recognise the low-pass filtered rhymes (e.g. children with DLD recognised 89% of the low-pass filtered rhymes compared to 42% of the bandpass filtered rhymes, while the TD children recognised 94% and 59% respectively). When relations with language development were investigated in the full sample of 9-year-olds (N = 95), accuracy in the bandpass-filtered speech condition was the stronger predictor of language outcomes, accounting for 12% of unique variance in the Clinical Evaluation of Language Fundamentals (CELF-III, Wiig, Semel & Secord, 2017) receptive language measures and 10% of unique variance in CELF expressive language scores. Accordingly, Goswami et al. (2016) suggested that although prior TSF studies of children with dyslexia and children with DLD indicated a shared sensory difficulty in processing slower temporal modulation patterns in speech, the neural temporal integration windows that are most impaired may differ between dyslexia and DLD. The speech data collected by Goswami et al. (2016) supported the RAP view that children with DLD may also have difficulties with faster temporal integration windows in speech-based tasks (gamma band difficulties).

Accordingly, we explored four *a priori* predictions in the current study, as well as conducting a series of analyses regarding potential group differences in cross-frequency neural dynamics. First, on the basis of the shared behavioural and sensory features of dyslexia and DLD and our prior infant data, we expected a different group preferred phase in the delta band for the children with DLD compared to the age-matched TD control children (H1). Second, on the basis of Araújo et al. (2024) and our TSF infant data, we tentatively predicted atypical delta-theta cross-frequency coupling for the children with DLD (H2), and greater theta-band power (H3). Finally, given the prior nonspeech neural data supporting RAP theory (Benasich et al., 2006, 2008; Cantiani et al., 2016, 2019; Choudhury & Benasich, 2011; Gou et al., 2011; Heim et al., 2011) and the filtered speech findings discussed above (Goswami et al., 2016), our fourth prediction concerned possible gamma band group differences (H4). Exploration of other cross-frequency dynamics (PAC between other bands, and phase-phase coupling [PPC]), ERPs and angular velocity were motivated in part by prior studies of children with dyslexia (Keshavarzi et al., 2022, 2024a). Prior data have been suggestive of different changes in phase across time for dyslexic children, but preserved PAC (Keshavarzi et al., 2024a). Accordingly, it was of interest to compare cross-frequency coupling variables and angular velocity for children with and without DLD.

## 2. Methods and materials

### 2.1. Participants

Sixteen typically developing children (mean age of 9.1 ± 1.1 years) and sixteen children with DLD (mean age of 9 ± 0.9 years) took part in the study. All participants were taking part in an ongoing study of auditory processing in DLD (see Parvez et al., 2024), and those children with suspected language difficulties were nominated by the special educational needs teachers in their schools. Children in the TD group were nominated by classroom teachers as being typically-developing. All children had English as the main language spoken at home. All participants exhibited normal hearing when tested with an audiometer. In a short hearing test across the frequency range 0.25 – 8 kHz (0.25, 0.5, 1, 2, 4, 8 kHz), all children were sensitive to sounds within the 20 dB HL range. Language status was ascertained by a trained speech and language therapist (the second author), who administered two subtests of the Clinical Evaluation of Language Fundamentals (CELF-V, Wiig, Semel & Secord, 2017) to all the participants, recalling sentences and formulating sentences. Those children who appeared to have language difficulties then received (depending on age) two further CELF subtests drawn from word structure, sentence comprehension, word classes and semantic relationships. Children who scored at least 1 S.D. below the mean (7 or less when the mean score is 10) on at least 2 of these 4 subtests were included in the DLD group. Oral language skills in the control children were thus measured using only two CELF tasks, recalling sentences and formulating sentences, and all control children scored in the normal range (achieving scores of 8 or above). Due to the Pandemic, although all 16 control children received the recalling sentences task, only 11 also received the formulating sentences task (all DLD children received 4 CELF tasks). The Picture Completion subtest of the Wechsler Intelligence Scale for Children (WISC IV, Wechsler, 2016) was used to assess non-verbal intelligence (NVIQ) and did not differ between groups. Group performance for the tasks administered to both groups is shown in Table 1.

**Table 1.** Details of the participating children, showing Scaled Scores (ScS, population mean = 10) and Standard Scores (SS, population mean = 100).

|  | DLD | Age-Matched Control (TD) |
| --- | --- | --- |
| N | 16 | 16 |
| Age (months) <sup>A</sup> | 108.3 (11.3) | 108.9 (13.3) |
| Recalling Sentences ScS <sup>B</sup> | 5.3 (1.7) | 10.9 (2.3) |
| Formulating Sentences ScS <sup>B</sup> | 4.4 (1.9) | 10.8 (1.7) |
| WISC Picture Completion ScS <sup>A</sup> | 7.9 (3.0) | 9.9 (2.8) |

Note. **A**: TD and DLD groups are not significantly different; **B**: DLD group significantly worse than TD group, $p < 0.01$ . WISC: Wechsler Intelligence Scale for Children.

### 2.2. Experimental set up and stimuli

The experimental set up and stimuli used in the EEG study were identical to those described by Keshavarzi et al. (2022, 2024a). After putting on the EEG cap, the children were seated in an electrically shielded soundproof room. They watched (on the screen) and listened (through earphones) to audio-visual stimuli. The EEG data were collected at a sampling rate of 1 kHz using a 128-channel EEG system (HydroCel Geodesic Sensor Net). The stimuli were a talking head presenting rhythmic sequences of the syllable “ba” at a rate of 2 Hz with both visual and additory information present. Visual cues began 68 ms before the onset of the stimulus “ba” as in natural speech. Each trial (sequence) consisted of a sequence of 14 repetitions of syllable “ba” with both visual and auditory information present (See Figure 1). One of the 14 syllables (either 9th, 10th or 11th syllable) in each trial could be out of time (rhythmic violation), in a random order. The temporal delay was calibrated individually and adaptively for each child, so that the oddball was detected around 79.4% of the time by a button press. The duration of each syllable “ba” was 280 ms, and the presentation level was 65 dB sound pressure level (SPL, measured using a Larson Davis 831 SPL meter). Each child was presented with 90 trials, 15 of which did not have a violation and were presented randomly as catch trials. Each trial contained three periods (see Figure 1): entrainment period, violation period, and return-to-isochrony period. Children were instructed to concentrate on the lips of the female face on the screen and to listen to the auditory stimuli. They were also asked to press a target key on the keyboard when one of the syllables was out of time. The total time for the EEG recording (excluding the set-up) was about 15 min. Via the adaptive procedure, the success rate in the behavioural task was equated for all participants. We calculated the temporal delay for both control (mean: 167.26 ms) and DLD (mean: 197.96 ms) children. We then applied a Two-sample t-test to compare the temporal delay between the two groups. The results suggested no significant difference (Two-sample t-test, *p* = 0.38).

**Figure 1.**
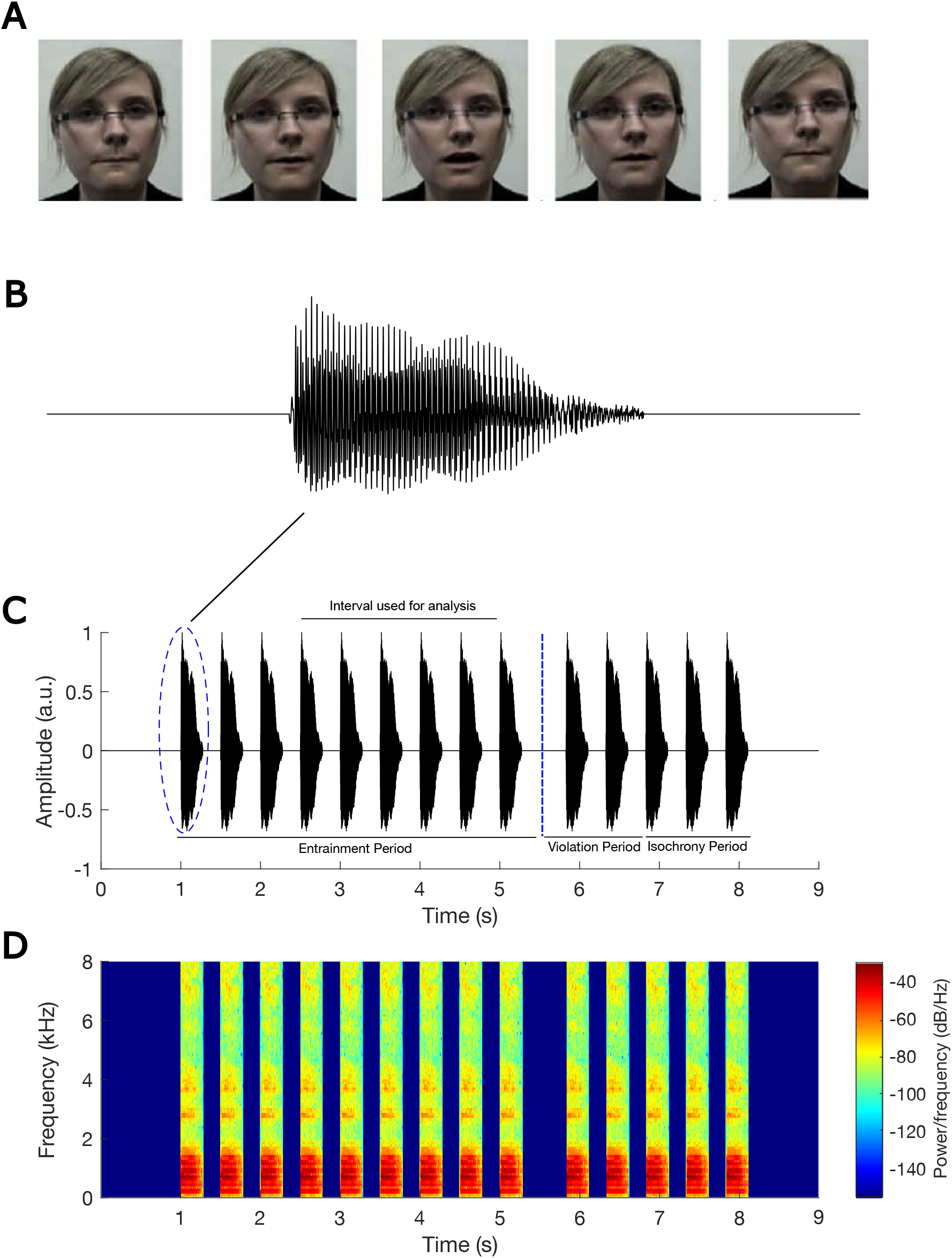
Experimental stimuli. Panel A shows frames of the visual information corresponding to a single syllable “ba”, panel B shows the waveform of a single syllable “ba”, panel C indicates an example trial (sequence of “ba” stimuli) with the violation at position of 10th stimulus, and panel D shows the spectrogram corresponding to sequence in panel C. This figure reproduced with permission from Keshavarzi et al., 2022.

### 2.3. EEG data pre-processing and analyses

The EEG data were first referenced to Cz and then band-passed filtered into 0.2 – 48 Hz using a zero phase FIR filter with low cutoff (−6 dB) of 0.1 Hz and high cutoff (−6 dB) of 48.1 Hz (EEGLab Toolbox; Delorme and Makeig, 2004). The bad channels were identified and interpolated using spherical interpolation (EEGLab Toolbox). The Independent Component Analysis (using FastICA algorithm, provided by EEGLab Toolbox) was run for each participant and the calculated independent components were then manually evaluated very carefully to remove artefactual components such as eye blinks and eye movements. This evaluation was based on the inspection of the scalp map, power spectrum, and time-domain waveform of the components. The pre-processed EEG data were downsampled to 100 Hz using function *resample*(). The data were either not filtered for power spectral analysis or band-pass filtered using function *filter*() into delta (0.5 – 4 Hz), theta (4 – 8 Hz) and low gamma band (25 – 40 Hz) frequency bands (MNE Library-Python, Gramfort et al., 2013) for measuring other metrics including phase entrainment, angular velocity, ERP, PAC and PPC. The data were then epoched into individual trials, which refers to the time interval starting 0.5 seconds before the onset of the first “ba” stimulus and ending 4 seconds after this onset. This interval encompasses the presentation of eight “ba” stimuli. In this study, the analyses were focused only on the EEG data corresponding to stimuli 4th-8th in the entrainment period of each trial to ensure that rhythmicity had occurred (Power et al., 2013).

### 2.4. Computation of angular velocity

The angular velocity, *w*(*t*), for signal *s*(*t*) over time interval Δ*t* = *t*_1_ − *t*_2_ is calculated as:

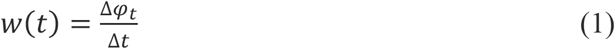

where Δ*φ*_*t*_ refers to the changes in the instantaneous phase (*φ*(*t*)) of the signal over the time interval Δ*t*. The instantaneous phase is calculated as:

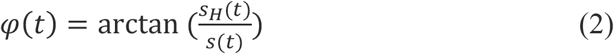

where *s*_*H*_(*t*) is the Hilbert transform of signal *s*(*t*), and is calculated as:

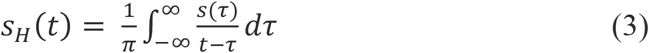

### 2.5 Computation of phase entrainment

To check the phase entrainment for each group in each frequency band, we performed the following steps (Keshavarzi et al., 2022):

1. Calculating instantaneous phases of all 128 EEG channels at the times corresponding to onsets of the 5 “ba” stimuli (4th-8th) in each trial.
2. Computing mean phase for each trial by averaging across the phase values obtained for all EEG channels and for the 5 “ba” stimuli in step 1. This resulted in a single-phase value for each trial.
3. Considering a single unit vector whose angle is determined by the phase value obtained in step 2 for each trial.
4. Calculating the mean vector for each child by averaging across the unit vectors in step 3. This resulted in a single vector for each child, called the *child resultant vector*.
5. Considering a single unit vector whose angle is determined by the phase of the *child resultant vector* in the vector space for each child.
6. Calculating the mean vector for each group by averaging across the unit vectors obtained in step 5. This results in a single vector, called the *group resultant vector*, for each group.

### 2.6. Computation of ERP

To obtain the time-domain ERP for each group and each frequency band, the following steps were conducted: (1) Calculating the mean voltage amplitude in the time interval of –500 ms to 500 ms relative to the individual “ba” (4th-8th “ba” in the trial) by averaging across the voltage amplitudes of all 128 EEG channels in the time interval; (2) Calculating the mean voltage amplitude for each trial by averaging over the mean voltage obtained (in step 1) for the 5 “ba” stimuli (4th-8th) in that trial; (3) Calculating the mean voltage amplitude for each participant by averaging across the mean voltage obtained (in step 2) for all trials of that participant; (4) Calculating the ERP for each group by averaging across the mean voltage obtained (in step 3) for all participants in that group.

### 2.7. Computation of the band-power

To investigate the band-power spectral density of the neural response over the time interval (including 4th-8th “ba”) used for analysis (See Figure 1C) for each each child and each group, the following steps were conducted: (1) Calculating the broad-band power spectral density of each EEG channel separately for each trial using the *periodogram*() function in MATLAB; (2) Calculating the broad-band power spectral density for each trial by averaging across the broad- band power spectral density obtained (in step 1) for channels; (3) Calculating the broad-band power spectral density for each participant by averaging across the broad-band power spectral density obtained (in step 2) for all trials of that participant; (4) Calculating the band-power spectral density for each frequency band (delta, theta, and low gamma) and for each participant based on the broad-band analysis obtained in step 3; (5) Calculating the band-power spectral density for each group and each frequency band by averaging across band-power spectral density obtained (in step 4) for all children in that group.

### 2.8. Computation of cross frequency PAC

Cross frequency PAC refers to the modulation of the amplitude of a signal at a high frequency band by the phase of low frequency oscillation. Here we quantified the strength of this type of modulation using the modulation index (*MI*; Tort et al. 2008; Hülsemann, 2019):

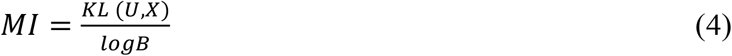

where *B* (=18) is the number of bins, *U* refers to the uniform distribution, *X* is the distribution of the data, and *KL* (*U, X*) is Kullback–Leibler distance which is calculated as:

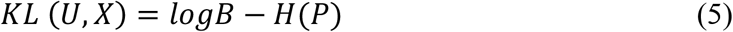

where *H*(·) is the Shannon entropy and *P* is the vector of normalized averaged amplitudes per phase bin which is calculated as:

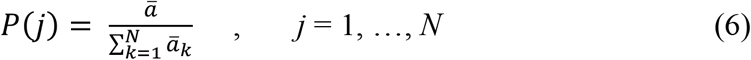

where *ā* is the average amplitude of each bin, *k* refers to running index for the bins. Note that *P* is a vector with *N* elements.

### 2.9. Computation of cross frequency PPC

Cross-frequency PPC refers to the phase synchrony between oscillations in two frequency bands. To quantify the PPC, we used the phase-locking value (*PLV*; Lachaux et al., 1999), which is calculated as:

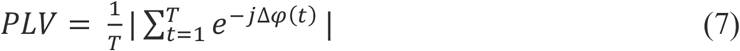

where |. | refers to the absolute value operator, *T* is the number of time samples, and Δ*φ*(*t*) is the phase difference, and is calculated as:

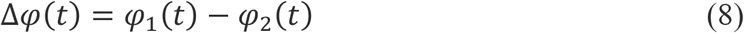

where *φ*_1_(*t*) is the instantaneous phase of first signal and *φ*_2_(*t*) is the instantaneous phase of the second signal.

## 3. Results

### 3.1. Phase entrainment consistency in each group

We expected consistent phase entrainment in each group in the delta and theta bands, but a different preferred phase in the delta band (H1). To assess phase entrainment consistency in each group and in each frequency band, we applied the Rayleigh test of uniformity to the child preferred phases separately for each group and for each frequency band (see Figure 2). For the control children, we found consistency in all bands: delta band (*z* = 5.14, *p* = 0.004, see Figure 2A), theta band (*z* = 3.70, *p* = 0.02, see Figure 2B), and low gamma band (*z* = 3.33, *p* = 0.03, see Figure 2C). For the children with DLD, we also found significantly consistent phase entrainment in the delta band (*z* = 8.31, *p* = 8.35 ×10^-5^, see Figure 2D) and in the theta band (*z* = 6.00, *p* = 0.002, see Figure 2E). However, phase consistency in the low gamma band was absent (*z* = 0.01, *p* = 0.99, see Figure 2F). This result indicates that the DLD brain does not show significant entrainment to rhythmic input in the gamma band, consistent with atypical neural responding to speech input in the low gamma band (H4).

**Figure 2.**
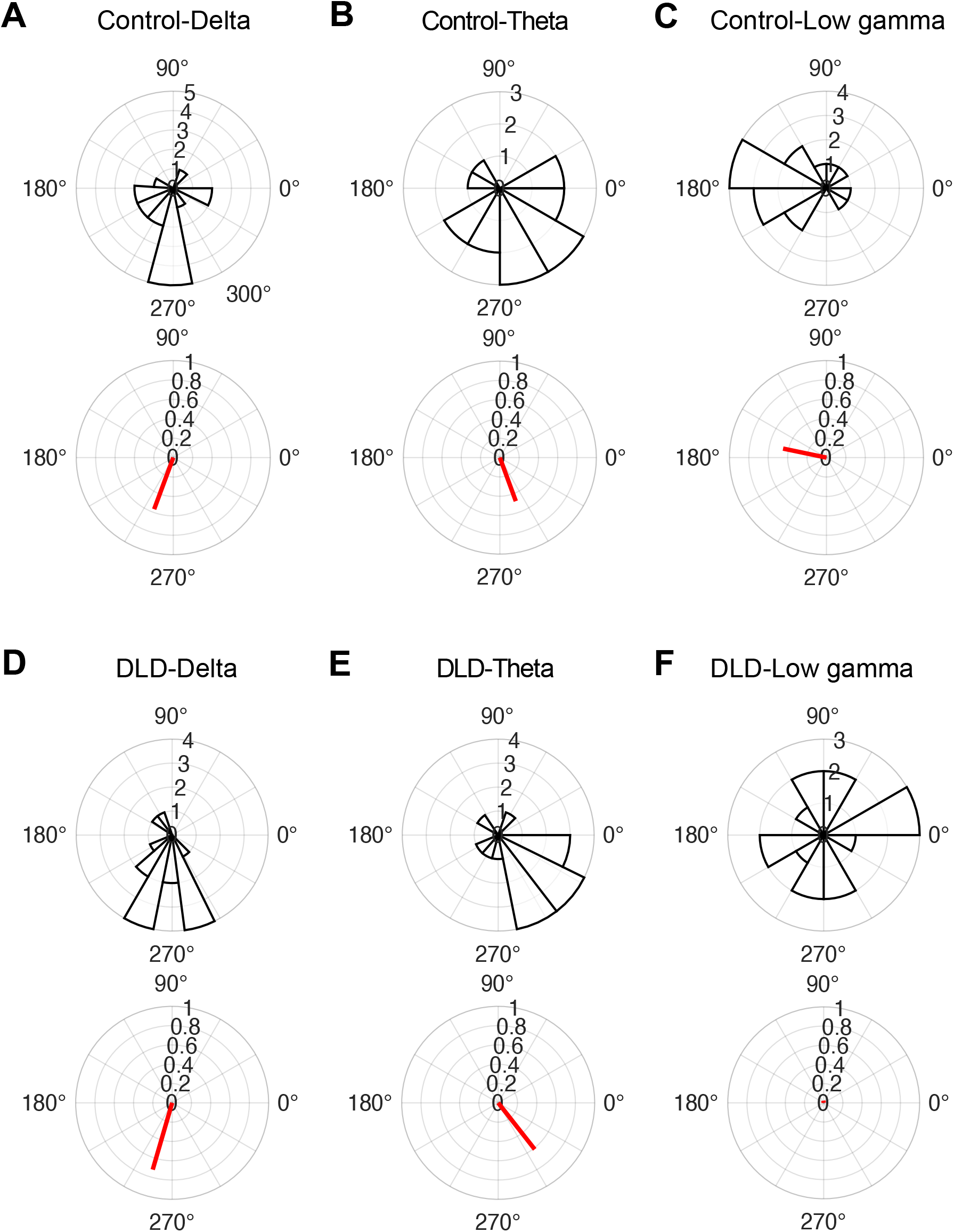
Entrainment phase distribution across children in each group (rose plots) and group resultant vectors (red lines). Note the absence of phase consistency in the low gamma band for the children with DLD.

### 3.2. Comparing the strength of phase entrainment between groups

The length of a *child resultant vector* can be considered as a criterion of the strength of phase entrainment across different trials for that child. Indeed, longer *child resultant vectors* indicate higher consistency of phase entrainment. To compare the strength of phase consistency in control children with children with DLD, we applied Wilcoxon rank sum tests based on the length of the *child resultant vectors* for the two frequency bands showing significant entrainment, delta and theta. There was no significant difference between the two groups for either band (delta: *z* = – 0.70, *p* = 0.49, Figure 3A; theta: *z* = – 0.55, *p* = 0.58, Figure 3B). We also checked these comparisons after removing outliers. A data point was considered as an outlier if the corresponding power value was greater than a 1.5 interquartile range above the upper quartile or below the lower quartile of the population data in that group. There were also no significant group differences after outlier removal.

**Figure 3.**
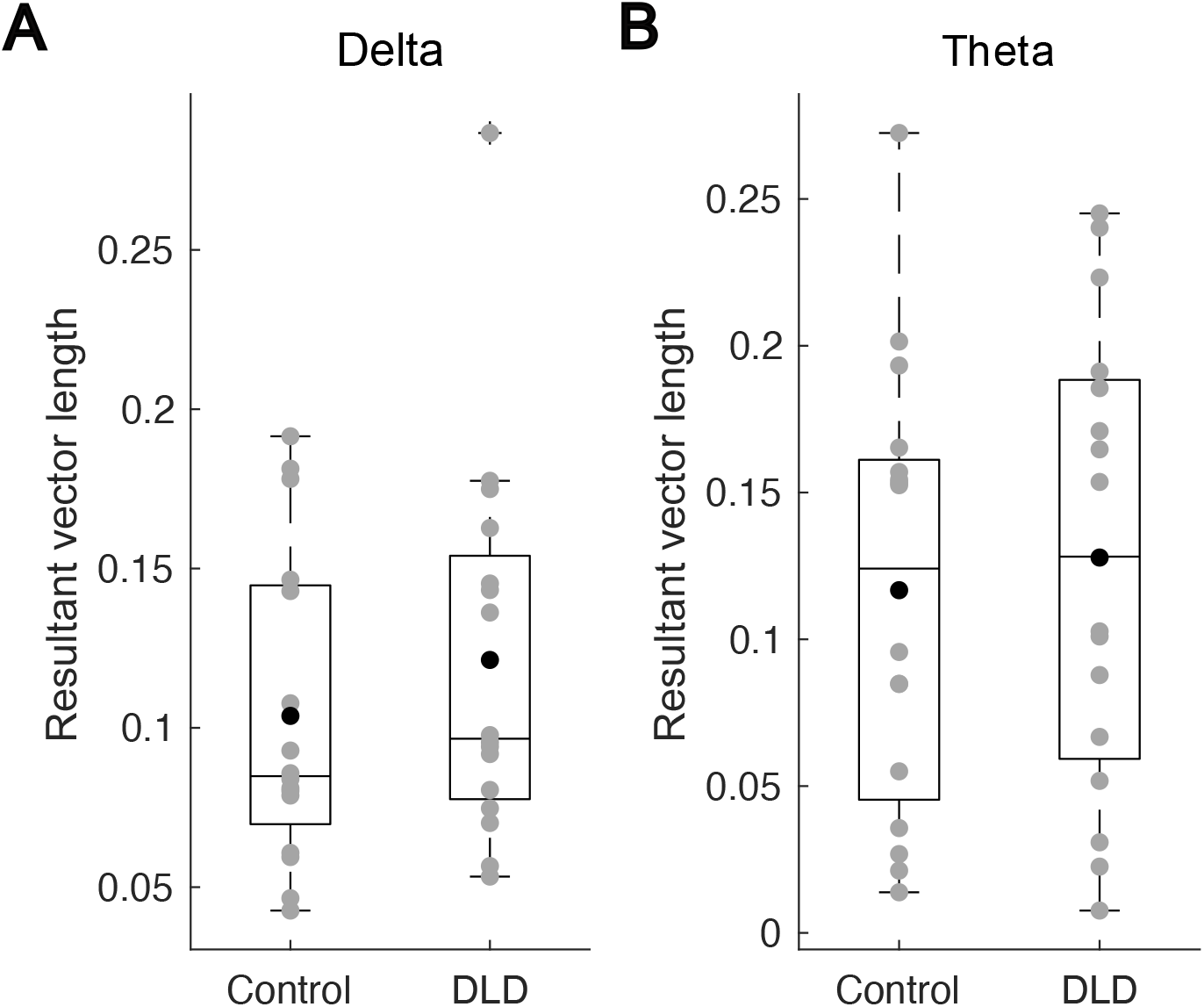
The length of child resultant vectors in the delta (A) and theta (B) bands in the two groups. The grey circles denote the length of the child resultant vectors for individual participants, the black disks denote the mean values for each group, and the horizontal line on each box shows the median.

### 3.3. Comparing preferred phase and angular velocity between groups

*A priori* we had expected group differences in preferred phase in the delta band (H1). To compare preferred phase between groups, the Watson-Williams test was applied to the delta and theta band data. The low gamma band had to be excluded, as it did not show consistent phase entrainment for the children with DLD. The Watson-Williams test assumes von Mises distributions, although this test is fairly robust against deviations from this assumption (Zar, 1999; Berens, 2009). In our data, the phase distribution for both delta and theta bands, as well as for both groups (TD and DLD), followed von Mises distributions (Watson’s U2 test, *p* > 0.05). Contrary to prediction, there was no significant difference in terms of preferred phase between the groups, neither in the delta band (Watson-Williams test, *p* = 0.84) nor in the theta band (Watson-Williams test, *p* = 0.46).

We also explored the behaviour of preferred phase across time separately for each group and each frequency band by computing angular velocities. Here the gamma band data could be included, as angular velocity (see Section 2.4) analyses do not require phase consistency, enabling a further test of H4. To this end, we calculated the group preferred phase for each frequency band prior to and after the occurrence of a stimulus (stimuli 4th–8th) in the entrainment period over the time interval of –500 ms to 500 ms. This time interval corresponds to two repetitions of the syllable “ba”. Figure 4 shows the group preferred phase for the two groups and for delta, theta, and low gamma bands. For both groups, delta-band and theta-band group preferred phases appear to follow a quasiperiodic pattern, with a frequency of around 2 Hz for the delta band (angular velocity of 4 rad/s, see Figure 4A) and a frequency of around 4 Hz for the theta band (angular velocity of 8 rad/s, see Figure 4B). For the low gamma band, however, the pre-stimulus group preferred phase for children with DLD appears to be different from that of the control children. We therefore employed a two-sample t-test to find the time bins with a statistical difference in terms of angular velocity. The result suggested a significant difference between the two groups over the time interval of –440 ms to –240 ms (two-sample t-test, *p* = 0.017, see Figure 4C). This suggests that phase changes differently across time for the children with DLD, in the gamma band only. The significant difference in angular velocity between the two groups during the pre-stimulus period (– 440 ms to – 240 ms) likely reflects differential anticipatory processes between DLD and the control groups. In this period, the brain is primarily involved in predicting the timing of the upcoming syllable, a process that might be more affected in children with DLD due to impairments in temporal prediction mechanisms. Once the stimulus is presented (post-stimulus period), the brain shifts from anticipatory to sensory processing. During this phase, children with DLD may exhibit more typical phase behaviour, which could explain the lack of significant differences in angular velocity from 60 ms to 260 ms. This temporal asymmetry in prediction versus processing may thus account for the observed pattern in our data.

**Figure 4.**
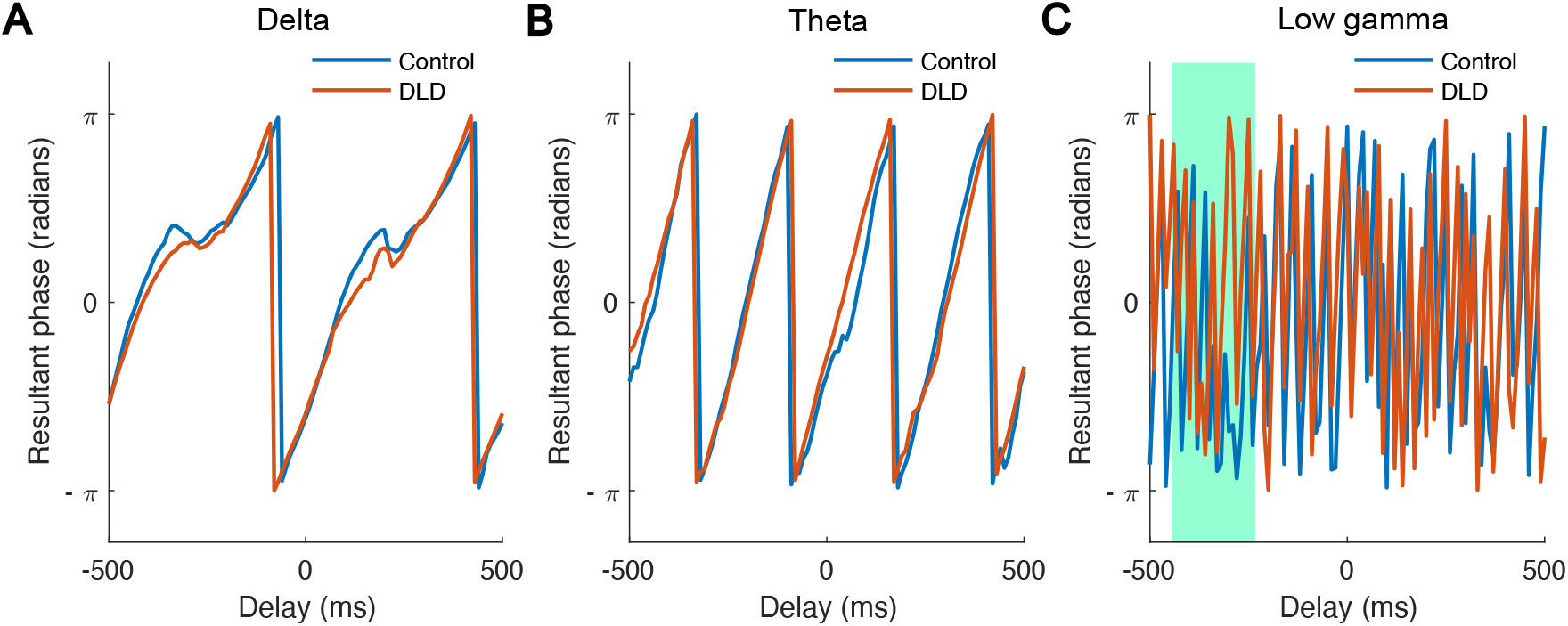
The group preferred phase (radians) as a function of the delay (ms) relative to the onset of “ba” stimuli. The blue and red curves are related to control (N = 16) and DLD group (N = 16), respectively. Both curves appear to follow a quasiperiodic pattern with a frequency of around 2 Hz in the delta band (A) and a quasiperiodic pattern with a frequency rate of around 4 Hz in the theta band (B). The curves related to the low gamma band (C) reveal a pre-stimulus difference between the two groups. In particular, the pre-stimulus angular velocity in this band over the time interval of –440 ms to –240 ms (marked with green colour) differs significantly by group.

### 3.4. Comparing ERPs between groups

Figure 5 depicts the ERP curves for each group. As we did not have an *a priori* prediction for ERPs, these analyses were exploratory in nature. We calculated the ERP in the time interval of –500 ms to 500 ms separately for each group and each frequency band, as described in Section 2.6. Although differences between group could be expected in terms of pre- and post-stimulus peaks in ERPs from visual inspection of Figure 5, no group difference reached significance in any frequency band (delta, theta, low gamma). Specifically, two-sample t-tests were conducted to compare the groups regarding the peaks (marked with green circles in the figure) in the time interval from –100 ms to 100 ms. The results indicated no significant group differences in delta (P1: *p* = 0.13, P2: *p* = 0.31), a trend for pre-stimulus differences in theta (P3: *p* = 0.053, P4: *p* = 0.26), and no differences for low gamma (P5: *p* = 0.30, P6: *p* = 0.78, P7: *p* = 0.19).

**Figure 5.**
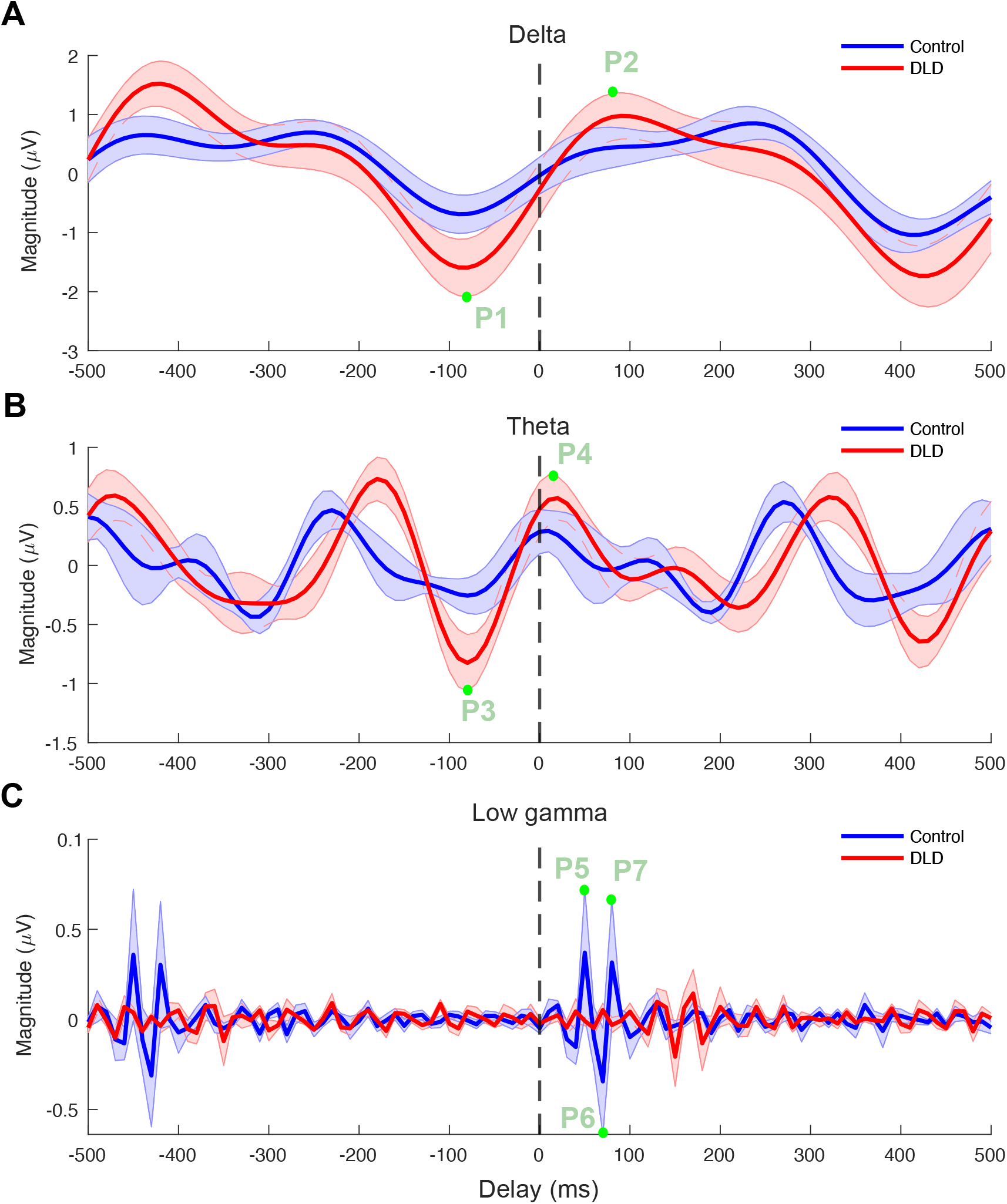
Event related potentials for delta (A), theta (B) and low gamma (C) bands. The blue and red curves denote the ERPs for control and DLD groups, respectively. The shaded areas denote the standard error of mean for each group. The ERP peaks compared statistically (P1 – P7) are marked with green dots.

### 3.5. Comparison of band-power between groups

We also conducted a broad-band power spectral analysis (as described in Section 2.7), with a particular focus on the 2 Hz stimulus presentation frequency and its harmonics (see Figure S1 in Supplementary Information). Despite rigorous preprocessing and spectral analysis, the expected peaks at 2 Hz were not clearly prominent, likely due to the inherent noise in EEG data collected from children. Indeed, low SNR can weaken the measured power at specific frequencies, particularly in populations such as children where movement artifacts, attention variability, and general signal instability can mask consistent oscillations that would occur under different conditions. To compare the band-power between groups, Wilcoxon rank sum tests were applied for each frequency band after removing outliers (3 control and 2 DLD outliers). In accordance with H3, we expected possible theta band group differences. We calculated the average power over the time interval used for analysis (see Figure 1C) in each frequency band separately for each group, using steps described in Section 2.7. Figure 6 shows the average power versus frequency separately for each group and for delta (see Figure 6A), theta (see Figure 6B), and low gamma (see Figure 6C) bands. A significant difference between control children and children with DLD was found in the theta (*z* = –2.11, *p* = 0.03; Figure 6B) and the low gamma (*z* = –2.69, *p* = 0.007; Figure 6C) bands. However, there was no difference in band-power in the delta band (*z* = –1.67, *p* = 0.09; Figure 6A). The effect sizes (Cohen’s d) for the delta, theta, and low gamma bands were calculated to be 0.74, 0.77, and 0.73, respectively, indicating medium to large effect sizes. This suggests that, despite individual variability, the observed differences are meaningful.

**Figure 6.**
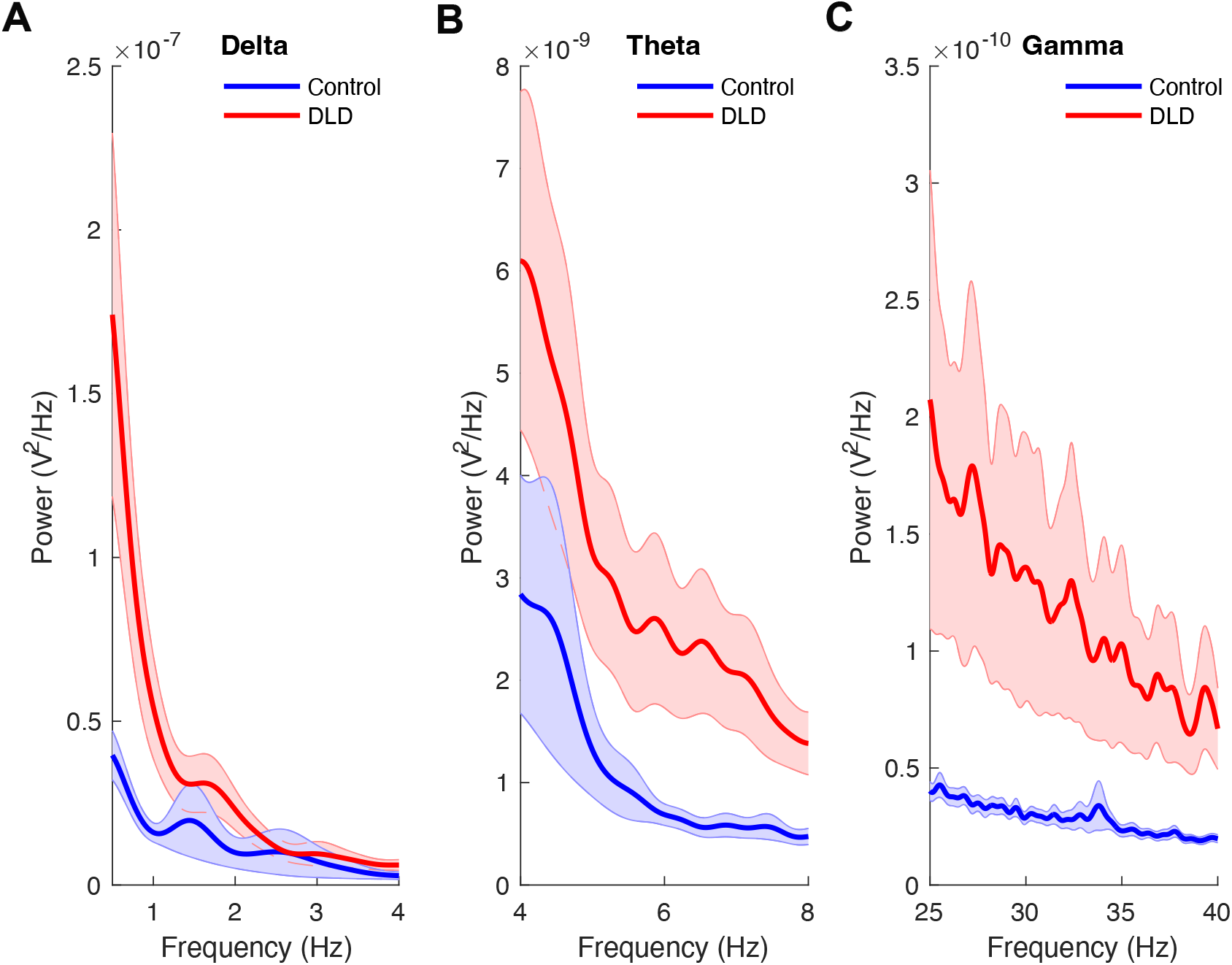
Delta-band (A), theta-band (B), and low gamma-band (C) power in the entrainment period. The shaded areas denote the standard error of the mean for each group. Please note the numerical power differences in the y axis by band.

### 3.6. Comparing cross-frequency PAC between groups

*A priori*, we had tentatively predicted potential group differences in delta-theta PAC between groups (H2). Delta-theta PAC, delta-low gamma PAC and theta-low gamma PAC were computed over the time interval used for analysis (see Figure 1C) separately for each group by computing the respective modulation indices. We then applied Wilcoxon rank sum tests to compare these modulation indices between the two groups. No significant differences were found for delta-theta PAC (*z* = –1.34, *p* = 0.18, Figure 7A) or delta-low gamma PAC (*z* = – 1.07; *p* = 0.28, Figure 7B), nor for theta-low gamma PAC (Wilcoxon rank sum test, *z* = –0.85; *p* = 0.40, Figure 7C). Similar non-significant results were found following outlier removal. Contrary to prediction, therefore, delta-theta PAC was not different in the children with DLD for the rhythmic syllable repetition task.

**Figure 7.**
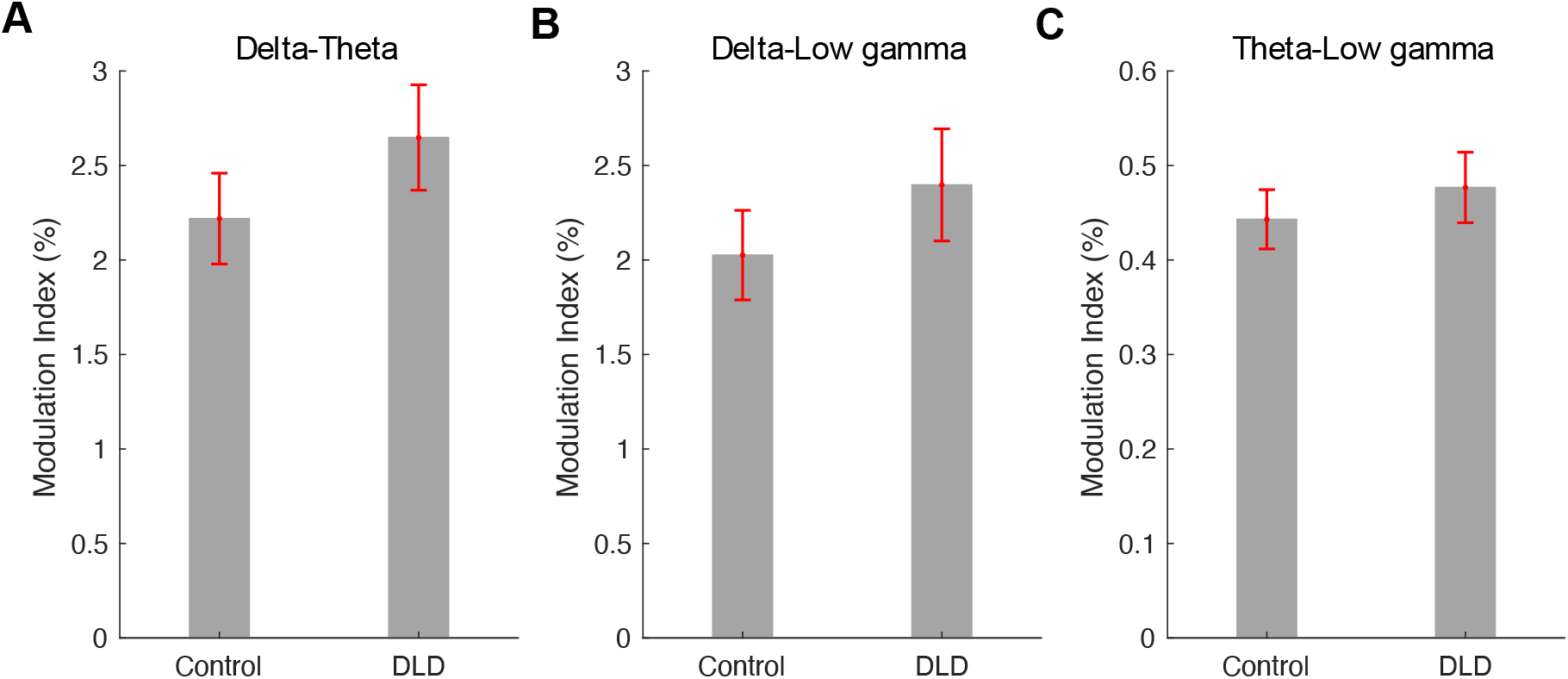
Modulation index was used as a measure for delta-theta (A), delta-low gamma (B), and theta-low gamma (C) phase-amplitude coupling. The error bars denote the standard error of mean.

### 3.7. Comparing cross-frequency PPC between groups

PPC was also compared between groups in further analyses. Delta-theta PPC, delta-low gamma PPC and theta-low gamma PPC were computed over the time interval used for analysis (see Figure 1C) separately for each group, as described in Section 2.9. We then applied two-sample t-tests to compare the two groups. As for PAC, no significant group differences were found, delta-theta PPC (*p* = 0.89, Figure 8A), delta-low gamma PPC (*p* = 0.26, Figure 8B), theta-low gamma PPC (*p* = 0.11, Figure 8C). Group PPC remained statistically equivalent after outlier removal.

**Figure 8.**
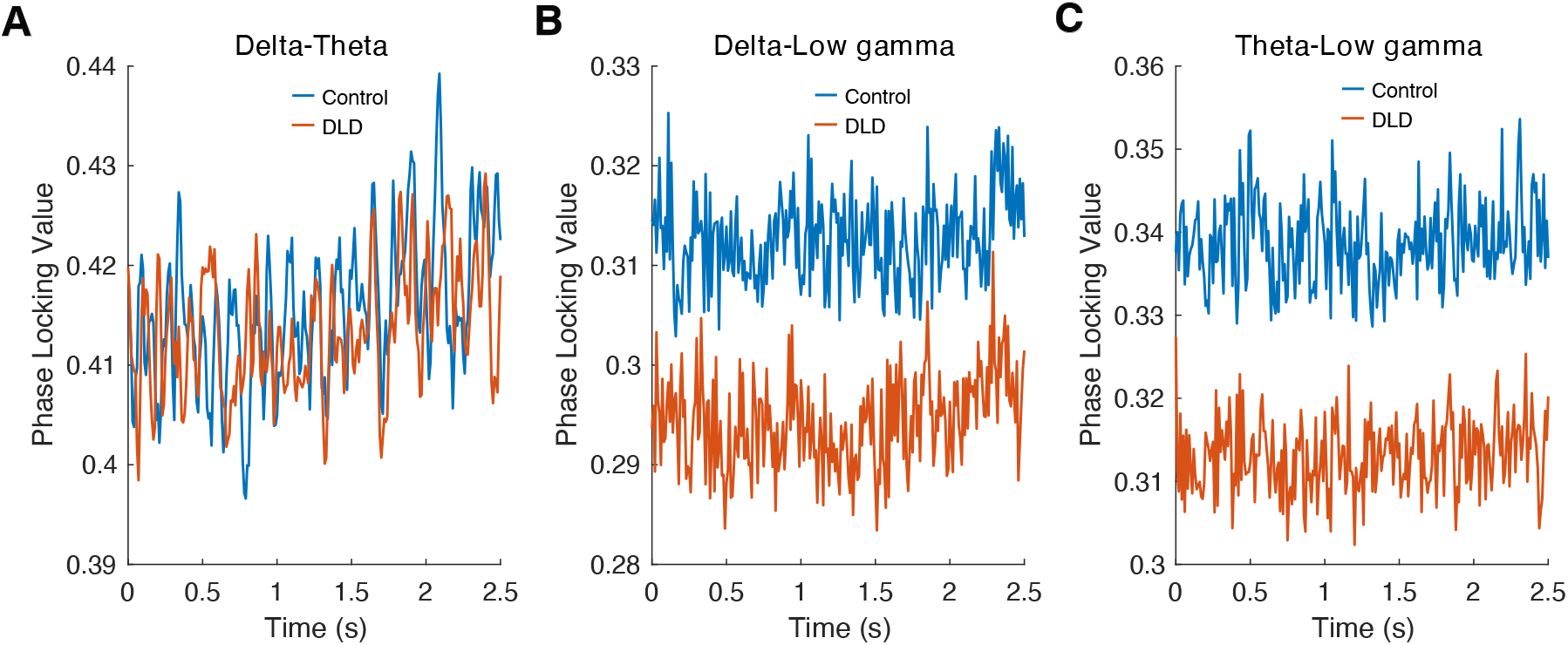
Delta-band (A), theta-band (B), and low gamma-band (C) phase-phase coupling. The blue and red curves denote the PPCs for control and DLD groups, respectively. Please note the different numerical scales on the y axis.

In the final analysis, we calculated the phase difference (see Equation 8) for delta-theta, delta-low gamma, and theta-low gamma oscillations for each child at the onsets of stimuli (4th–8th) during the entrainment period. The resultant phase difference for each group was then computed by averaging across the phase differences obtained for children in that group, and separately for delta-theta, delta-low gamma, and theta-low gamma oscillations (see Figure 9). We applied Watson-Williams tests to compare the group resultant phases between the groups. All measures for both groups (TD and DLD) followed von Mises distributions (Watson’s U2 test, *p* > 0.05). In this case, low-frequency resultant phase (delta and theta) did show significant differences between the two groups when coupling with the higher-frequency gamma band, delta-low gamma (*p* = 0.04), theta-low gamma (*p* = 3.28 × 10^-4^). Accordingly, the synchronisation of low-frequency phases (delta, theta) with high-frequency phase (low gamma) appears to operate differently for children with DLD, providing some support for H2. These differences are clearly visible in Figure 9. However, regarding delta-theta resultant phase differences, the two groups were not significantly different (*p* = 0.11).

**Figure 9.**
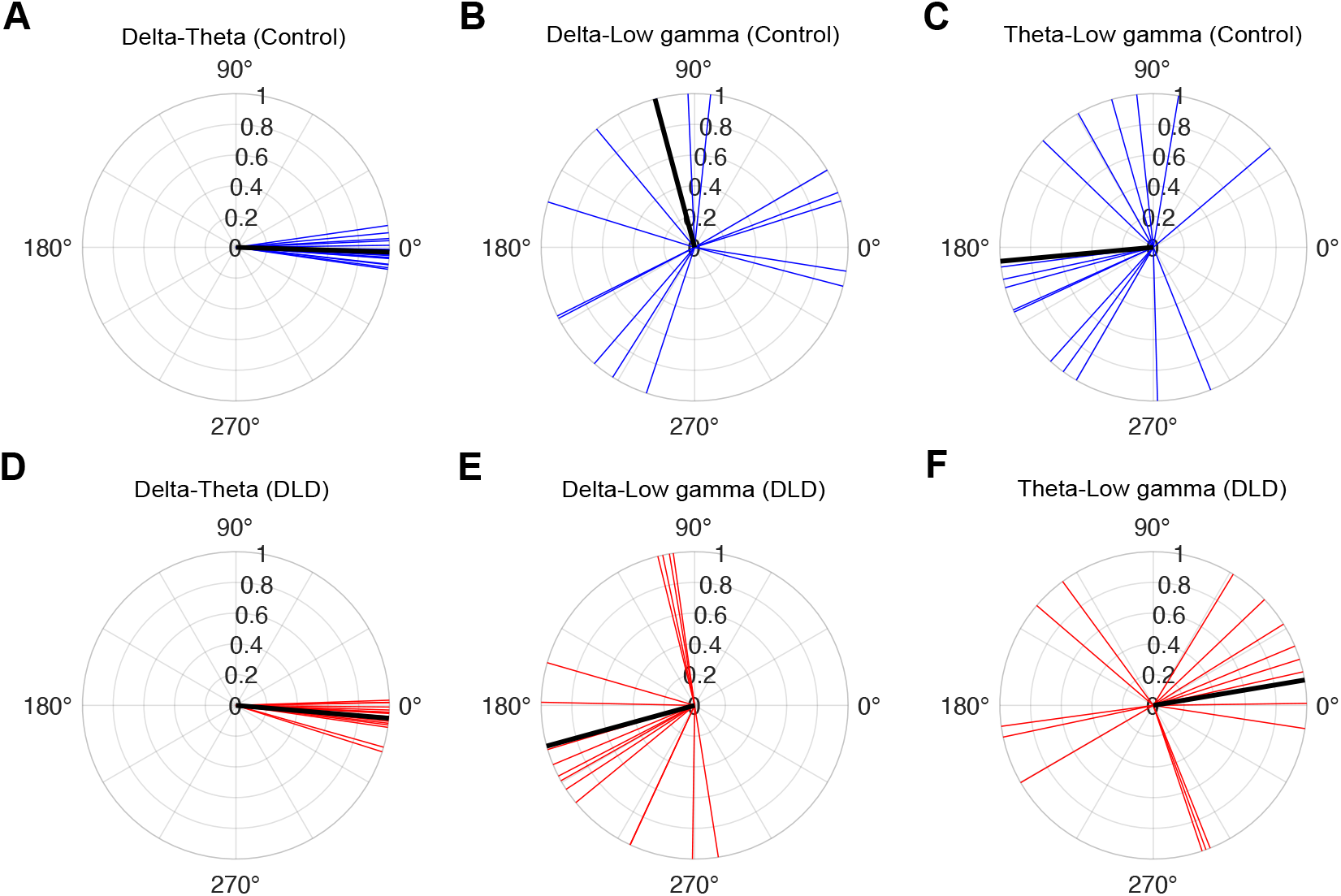
Delta-theta, delta-low gamma, and theta-low gamma resultant phase differences at the onsets of stimuli (4th–8th) during the entrainment period for control (A, B, C) and DLD (D, E, F) children. The blue and red vectors denote the resultant phase difference for individual children in control and DLD groups, respectively. The black vectors show the group resultant phase difference.

## 4. Discussion

The current study investigated the processing of speech rhythm by children with DLD at the neural level, by examining different mechanistic aspects of the oscillatory encoding of a highly constrained rhythmic speech input, the repetition of the syllable “ba” at a 2 Hz rate. Four *a priori* hypotheses were investigated. First, given the similar sensory and cognitive profiles of children with DLD and children with dyslexia in behavioural speech rhythm tasks, we expected that children with DLD would show a different preferred phase in the delta band compared to typically-developing (TD) age-matched control children (H1). Second, we predicted that dynamic relations between the delta and theta bands may be different for children with DLD when compared to TD control children (H2). Third, we expected to find greater theta power during rhythmic speech processing in the children with DLD (H3). Finally, we expected that gamma band processing may be atypical in the children with DLD (H4).

Regarding H1, the expectation of impaired delta phase for the children with DLD was not supported. The children with DLD showed equivalent phase entrainment strength to TD children in the delta band, and they also showed equivalent preferred delta phase. Further, they showed equivalent delta band angular velocity, and ERPs in the delta band did not differ by group either. Delta band power in the entrainment period was equivalent by group. Accordingly, despite the similarities in behavioural and sensory rhythm processing tasks that have been documented for children with DLD and children with dyslexia (Richardson et al., 2004; Corriveau et al., 2007; Thomson & Goswami, 2008; Corriveau & Goswami, 2009; Fraser et al., 2010; Goswami et al., 2011; Beattie & Manis, 2012; Goswami et al., 2013; Cumming et al., 2015a,b; Richards & Goswami, 2015, 2019), the neural mechanisms that are atypical for children with DLD in this paradigm were quite different from those previously documented for children with dyslexia (Power et al., 2013; Keshavarzi et al., 2022), at least for the current participants.

Regarding H2, cross-frequency coupling between delta and theta band responses was not significantly different between the DLD and TD children. The analyses of cross-frequency dynamics (PAC, PPC, and angular velocity) did not reveal group differences when delta and theta band responses were measured (Figures 7 and 8). Visual inspection of Figure 7 shows that delta-theta PAC had a higher (though non-significant) modulation index for the children with DLD, a result that was also found by Araújo et al. (2024) with a smaller DLD group (for whom it was significant). Regarding H3, theta band power during the entrainment period was significantly greater in DLD children (Figure 6). Further, when group differences in resultant phase were computed in exploratory analyses, theta-low gamma resultant phase differed significantly between groups, as did delta-low gamma phase. Hence when low-frequency phases had to synchronise with high-frequency gamma responses in our task, there were group differences (Figure 9).

Consistent with H4, low gamma band oscillatory responses in the rhythmic syllable repetition task showed a number of differences in the DLD brain. Low gamma band phase entrainment in the children with DLD was absent in the current paradigm, suggesting that phase consistency in this band was not attained, despite the highly predictable input. Additionaly, children with DLD exhibited significantly greater low gamma power compared to the control children. There was also a significantly faster angular velocity in the gamma band for the DLD children, suggesting that gamma phase changed differently over time when compared to TD children. Furthermore, as noted, in the resultant phase analyses both delta-low gamma and theta-low gamma coupling was atypical (Figure 9). The resultant phase differences suggest a different underlying relationship between low-frequency phase and high-frequency phase in the DLD brain. One way of interpreting the consequences for speech processing is that the interactions or connectivity between neural networks at these different rates is impaired, namely the theoretical notion of *communication through coherence* (Fries, 2015). Fries (2009, 2015) proposed that neuronal inputs that arrive at random phases of the excitability cycle of related rhythmic neuronal responses will have lower effective connectivity, with consequently negative effects for cognition. Fries used examples from gamma band activity during visual cognition (primarily in animals) to illustrate communication through coherence, but the fundamental principle that he was proposing is relevant here, namely that the synchronization of brain oscillations is a key mechanism for information encoding and perceptual binding.

In our data, the gamma rhythmic response to the highly predictable speech input does not attain phase consistency, making it more difficult for low-frequency responses (delta, theta) to synchronise their phases to gamma phase. This could suggest reduced coherence during speech processing, negatively affecting the precision of linguistic encoding. In communication through coherence theory, gamma-band coherence enables communication to be not only effective but also precise. Although some aspects of our data suggest that the low-frequency responses (delta and theta) and their phase relations are operating in the same way as for TD children during the rhythmic speech task (with equivalent PAC and PPC strength), the data also suggest that an optimal phase relation with gamma band responses in the temporal domain cannot be established, thereby impairing speech processing. This is mechanistically quite different from our previous findings with children with developmental dyslexia in this paradigm, where delta phase itself was atypical (Power et al., 2013; Keshavarzi et al., 2022). Nevertheless, for dyslexic children also, PAC mechanisms were explored and appear to be intact (delta-beta PAC, see Power et al., 2016; Keshavarzi et al., 2024a).

A related view of the importance of gamma band processing for speech encoding concerns the parsing of phonemic information in the speech signal (Giraud & Poeppel, 2012; Poeppel, 2014). Indeed, the RAP theory of DLD predicts impaired phoneme representation in affected children, as speech information in temporal integration windows of ∼40ms is thought to arrive too fast for efficient processing by children with DLD (Tallal & Piecey, 1973; Tallal, 1980). Our data is partially consistent with RAP theory, as resultant phase mechanisms may contribute to impaired phonemic processing. However the temporal integration window in RAP theory is based on non-speech tone tasks. Regarding oscillatory *speech* encoding mechanisms, the phoneme parsing view is based on the temporal rates thought to be required rather than on empirical data (Giraud & Poeppel, 2012). However, a recent French study with dyslexic adults reported that 20 minutes of tACS stimulation at ∼30Hz significantly improved phonemic awareness in a behavioural task (Marchesotti et al., 2020). The data presented here may suggest that the ∼30 Hz response to acoustic feature information in speech is atypical in the DLD brain, which could impair some aspects of phoneme awareness. Nevertheless, most phonetic features are present in natural speech over much longer time windows than 40 ms (see Menn et al., 2023). Further, the cortex maintains representations of phonetic information even after it is lost from sensory input (Gwilliams et al., 2022). Accordingly, the idea that the gamma band differences reported here prevent children with DLD from representing phonemic information per se seems unlikely. Accordingly, we prefer to interpret the data using communication through coherence theory. Our findings appear to indicate reduced co-ordination of low-frequency oscillatory phase with gamma band responses in the DLD brain, reflecting altered neuronal network interactions during linguistic processing that may affect encoding precision and thereby impair speech perception at all phonological levels, not only the phoneme level. Converging studies with natural speech rather than the rhythmic speech paradigm utilised here are required to assess these different possibilities.

The current study has a number of limitations. Firstly, the rhythmic speech task is not a natural language task. Further investigation of the impaired rhythm hypothesis using natural speech listening tasks is required in order to establish whether the current gamma band and resultant phase differences are present during natural language processing. Second, the sample size is relatively small, with 16 DLD children. However, it is representative of the existing neural DLD literature (e.g., Heim et al. 2011 studied 17 children with DLD, Choudhury & Benasich 2011 studied 17 infants at risk for DLD, Nora et al. (2024) studied 17 children with DLD). This reflects the difficulty of engaging families who have children with DLD for neuroimaging studies. Thirdly, the presentation format that we used did not enable separation of visual entrainment mechanisms from auditory-visual mechanisms, due to time constraints in testing. This separation has been achieved in our prior studies of both dyslexia (Power et al., 2013) and TD infants (Ni Choisdealbha et al., 2023). It is possible that visual speech cues play a key developmental role in temporal prediction in DLD, hence in future studies it would be important to measure the potential role of visual speech mechanisms also.

In conclusion, here we find that the 9-year-old DLD brain does not show the delta-band deficits that are predicted by TS theory to underpin impaired rhythmic speech processing. Instead, the children with DLD exhibited an absence of phase consistency in the gamma band when processing highly predictable rhythmic speech, and showed different angular velocity in the gamma band compared to TD children. The children with DLD also showed significantly greater gamma and theta power during the task, and a difference in resultant phase when low-frequency phases (delta, theta) coupled with low gamma. Overall, the data suggest reduced coherence in the DLD brain between slower and faster networks during speech processing, which would negatively affect the precision of linguistic encoding. Further studies employing natural continuous speech are required to investigate this interpretation in more depth.

## Data and Code Availability

Data and code will be made available on request.

## Author contributions

M.K. conceptualisation, methodology, data collection, data analyses, visualisation, writing – original draft; S.R. data collection, writing – review & editing; G.F. data collection, writing – review & editing; L.P. data collection, writing – review & editing; U.G. funding acquisition, project administration, supervision, conceptualisation, methodology, writing – original draft.

## Acknowledgements

The authors would like to thank all the children, families and schools involved in the study. This research was funded by a donation to UG from the Yidan Prize Foundation. The sponsor played no role in the study design, data interpretation nor writing of the report.

