## Supplementary Information for "Neural processing of rhythmic speech by children with developmental language disorder (DLD): An EEG study"

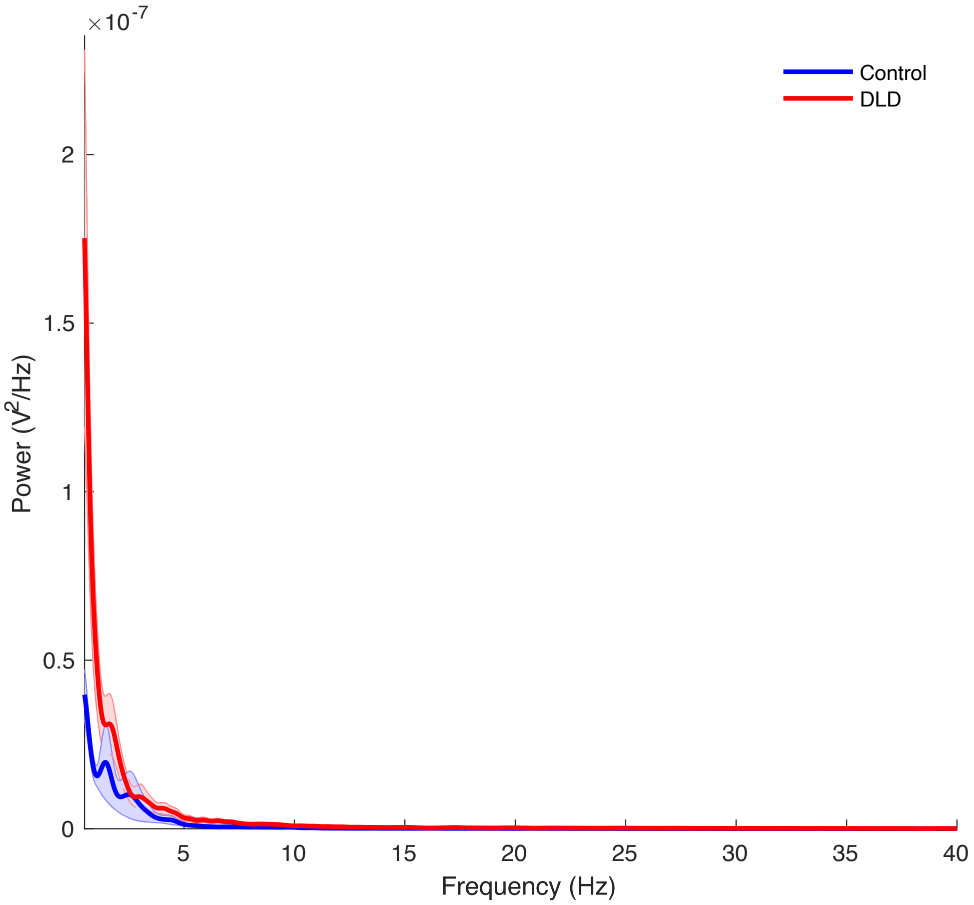


**Figure S1. Broad-band spectral power in the entrainment period.** The shaded areas denote the standard error of the mean for each group.
